# Plasmids from a complex biome exist as communities

**DOI:** 10.1101/2024.05.02.592190

**Authors:** Cian Smyth, Robert J Leigh, Thi Thuy Do, Fiona Walsh

## Abstract

Plasmids play a crucial role in the spread of antimicrobial resistance genes (ARGs) across One Health due to their ability to transfer a wide range of ARGs within and across bacterial species and biomes. We sequenced 173 circularised plasmids transferred from wastewater treatment plant (WWTP) effluent into *Escherichia coli* and subsequently characterised their genetic content. Multiple multidrug resistant plasmids were identified with a significant number of mega plasmids (>100Kb). Plasmids existing in isolation were rare and almost all existed with other plasmids. Our results suggest that positive epistasis promotes plasmid persistence in WWTP populations in a similar manner to that identified *in vitro* via infectious transmission, varying properties against plasmid community backgrounds, interactions with a range of other plasmids, source-sink spill-over transmission within the plasmid community rather than the host bacteria and compensatory mutations. We have demonstrated that the plasmid paradox solutions apply to plasmid communities in addition to plasmid host interactions. Our study identified that rather than existing as lone entities plasmids co-exist in small packs, the protection is afforded to the pack not by all members but by one or two and many plasmids coast within this pack as they contain no obvious advantage to the host. Our findings show that we need to enter a new paradigm and study plasmids in packs rather than as single entities in order to understand their transmission across One Health.

## Introduction

Antimicrobial resistance (AMR) is a global problem with the movement and transfer of AMR across One Health a concern for the emergence of novel resistance mechanisms, spread of known mechanisms and the increase in abundance of AMR genes (ARGs). We need to determine what mechanisms drive AMR plasmid emergence, spread and persistence within and across biomes to understand and predict their ecology (1). We must first characterise the plasmid communities in nature and within bacterial hosts capable of traversing biomes e.g. *Escherichia coli*.

Data on plasmid maintenance and their survival within bacterial populations is provided using infection principles from Stuart and Levin from the 1970s, which incorporated plasmid stability and plasmid maintenance (2). The introduction of co-existing or co-infecting plasmids was analysed in terms of plasmid persistence of the plasmid containing the selective advantage e.g. ARGs. However, plasmids were not described as co-existing communities of plasmids and the experiments defining these findings used artificially generated co-existing plasmids within the bacteria or specific examples from clinical pathogens (3, 4). Plasmids are not usually studied as semi-independent evolving entities like viruses. A recent study highlighted the evolutionary biases when observing plasmids in such a lens (5). However, the understanding we have of how plasmids interact, how they are selected, their fitness costs and their likelihood of transfer and persistence is based mainly on plasmids analysed *in vitro* and in the most part plasmids isolated from human pathogens as single entities (3).

The plasmid paradox suggested that where costs are associated with the plasmid they should be lost and where benefits are accrued from the plasmid they should be integrated into the chromosome, this paradox has been solved to explain why plasmids exist using ecology and evolution to explain their existence (6). However, the paradox theory presumes that plasmids have a cost of maintenance, which was not yet measured in the real world. Solving this paradox presents the idea that “Co-infecting plasmids alter the fitness effects (decrease) of plasmid acquisition, potentially masking fitness costs through epistasis” (6). However, this suggests that the function of co-infecting or co-existing plasmids is to ameliorate the effects of the plasmid with the selective gene(s) and the cost associated with the maintenance of these plasmids, which ignores the alternative roles of the co-existing plasmid or that they may co-exist in nature for other benefits or for the benefit of the plasmid lacking the selective advantage. In addition, the added complexity of mobilizable plasmids that rely on conjugative plasmids for their transmission further complicates co-existing plasmid dynamics, persistence and co-survival.

Scientists have artificially co-infected bacteria with two different plasmids to understand co-infection dynamics (3). While this results in the description of the two plasmids (one large and one small) tested it does not describe co-infection or co-existence of two plasmids from a community of plasmids that exist in nature. The importance of the variation is that not all plasmids can exist in the same bacterial cell due to size and the selection of a co-existing partner plasmid is an important step in the selection, ecology and evolution of plasmids within bacterial communities in the real world.

This study analysed antimicrobial resistance plasmid ecology at three levels:

1. Plasmid populations from the real world of WWTP effluent
2. Plasmids as co-existing communities within the bacterial host
3. Plasmid to plasmid variation under the same selection pressure and within the same bacterial host

Incompatibility groups frequently prevent the co-existence of plasmids in the same isolate, which is well proven for conjugative plasmids. Exclusion factors are also important but are less widely studied. The capacity for plasmids to co-exist is an important factor frequently overlooked in the analysis of plasmid dynamics, persistence and selection as it enables plasmids to “hide”. This adds complexity to the dynamics and selection of plasmids which has been rarely studied and even less so (if not at all) in a complex environment. The community context of the bacteria of the microbiome hosting the plasmids will impact the persistence and transmission of plasmids (7). While the host range of a plasmid will determine how far and wide the plasmid may spread its ability to co-exist with other plasmids will also determine how well it can hide under selective pressure for which it does not contain a gene or how it can persist within a population.

## Methods

### Sample Collection

Effluent collected form two WWTPs were used as the source of plasmids to be transferred into *Escherichia coli* (8). Exogenously extracted plasmids were transferred to *E. coli* from WWTP effluent samples (n=43) using previously described methods (9). The resulting *E. coli* with plasmids were selected on ampicillin or tetracycline or colistin at CLSI guidelines breakpoint concentrations (10).

### Plasmid Extraction

Plasmid extractions were performed using the Macherey-Nagel NucleoSpin Plasmid kit following the low-copy number protocol according to the manufacturer’s guidelines.

### Short-Read Sequencing

Illumina short-read sequencing was initiated with plasmid extractions quality assessment using NanoDrop and Qubit as per the sequencing centre (Novogene) guidelines. Extracted DNA was sequenced by Novogene using an Illumina NovaSeq 6000 with PE150 and Q30 ≥80[%. This provided far greater than 100X coverage for each plasmid.

### Long-Read Sequencing

Long-read sequencing was performed on all extracted plasmids using the Oxford Nanopore Technologies (ONT) MinION. Ligation library preparation was performed using the SQK-LSK-109 Ligation Sequencing kit with no noticeable deviation from protocols. Multiplexing was performed with the NBD-104 Barcoding kit.

### Pre-assembly Quality Control (QC)

Raw short reads were filtered and trimmed using Cutadapt *v*.3.0 (11). Raw long reads were processed using Filtlong *v*.2.0 for size and quality, with demultiplexing steps and adapter removal utilising Guppy v.6.1.2 (12, 13).

### Hybrid-Assembly

Hybrid assembly was performed using Unicycler *v*.0.5.0 with default settings (14).

### Post-assembly QC

Visual assessment of assembled contigs was performed with Bandage *v*.0.8.1 (15). This allowed for an easy inspection of circularised contigs.

### Plasmid Identification and Annotation

Plasmid identification among consensus sequences was performed using Platon v.1.5.0 utilising default settings (16). Each circularised plasmid was annotated using BAKTA *v*.1.2.2 using default settings.

### Antimicrobial resistance, metal resistance, virulence genes and replicon profiling

Each circularised plasmid was analysed for the presence of ARGs using ABRicate *v*.1.0.1 for AMR using the Comprehensive Antimicrobial Resistance Database (CARD) *v*.3.09 (17, 18). Each circular plasmid was profiled for metal and biocide resistance with BacMet *v*.2.0 (using a previously published back translated dataset) and for virulence factors using the Virulence Factor Database (VFDB) *v*.0.5, plasmid replicon type was determined using PlasmidFinder *v*.2.1 (19–22). As ABRicate requires a nucleotide input and as BACMET is only provided in amino acid format, the amino acid database was back translated (using translation table 11) prior to annotation (Supplementary information).

### Mobilisation Determination

Predicted plasmid mobilisation was determined using the MOBSuite, MOBTyper *v*3.0.3 with default settings (Supplementary information) (23).

### Plasmid rearrangement analysis

The first clustering of plasmids found them to be in a small number of groups using rounds of sequence comparisons using NUCmer (24). The data was then reformatted to generate the coordinates on both plasmids using the NucDiff wrapper around NUCmer (25).

## Results

The presence of ARGs in WWTP effluent has been well documented. However, the presence of these ARGs on plasmids and the characterisation of such plasmids is less well understood. The WWTP effluent samples analysed in this study had previously been analysed for ARGs using qPCR arrays, antibiotic chemical composition and microbiome content (8, 26, 27). There was a trimodal distribution of plasmid size which delineated the plasmids into small 6314bp or less, medium size of 34Kb to 55Kb and large of 96Kb to 292Kb detected in this study. The plasmids were also delineated into two groups: AMR plasmids and non-AMR plasmids. Communities of two or more plasmids predominated and single plasmid occupied *E. coli* were rare. The most clinically relevant ARGs were those conferring resistance to colistin or the fluoroquinolones.

### Antimicrobial resistance plasmids

Plasmid mediated AMR genes (n = 33 different genes) conferred resistance across ten antimicrobial classes (aminoglycosides, quinolones/fluoroquinolones, macrolides, polymyxins, beta-lactams, rifamycins, chloramphenicols, dihydrofolate synthesis inhibitors, sulfonamides and tetracyclines) (Supplementary information). Plasmids frequently carried seven ARGs and up to 12 ARGs were detected on one plasmid.

Colistin resistance gene *mcr-9* was detected on highly related plasmids all 287Kb or 292Kb. These plasmids also contained aminoglycoside, beta-lactam (*bla*_TEM-1_), ESBL (*bla*_SHV-134_), trimethoprim, tetracycline, macrolide and sulphonamide resistance genes. The 292Kb *mcr-9* plasmids in addition contained the quinolone resistance gene *qnrA* and two copies of *sul1*. These plasmids contained the replicon groups IncHI2A, IncHI2 and RepA. The *mcr-9* gene was flanked by and IS5-like element IS903B and an IS6-like element IS26. The 287Kb plasmid was highly similar to the 292Kb plasmids. However, in the 287Kb plasmids the *ampR*, *qnrA* gene and one copy of *sul1* was absent due to an absence of a 3147bp section of plasmid. The *mcr-9* plasmids also contained genes conferring resistance to arsenic, mercury, nickel, lead, copper, potassium tellurite and quaternary ammonium compounds (Supplementary information).

Plasmids of 54 - 55Kb size and containing the IncN replicon group contained both plasmid mediated quinolone resistance (PMQR) genes *aac(6’)Ib-cr* and *qnrB* genes and genes conferring resistance to aminoglycosides (*aadA16*), trimethoprim (*dfrA*), sulphonamide (two copies of *sul1*) and tetracycline (*tetA*) (Supplementary information). All resistance genes, except *tetA*, were contained together on a class 1 integron. The only metal resistance gene was *qacEdelta1* and there were no virulence genes.

### Plasmids with no resistance or virulence genes

The plasmids which contained no AMR, metal resistance nor virulence genes varied in size from 2337bp to 113,760bp (Supplementary information). The 32A plasmid was the largest at 113,760 bp, a non-mobilizable plasmid containing three phage types. This was essentially a plasmid of phage and contained an IncFIB replicon. Within this group of plasmids only the 56Kb was a conjugative plasmid, which contained the IncX replicon. The remainder of the plasmids above 7,000bp were approximately 34Kb. The 34Kb plasmids were IncP, and mobilizable but non-conjugative. They all contained heat resistance genes, which may serve as the selective pressure for their persistence. Most of the plasmids less than 7Kb contained a Col replicon but some contained no replicon. Most of these small plasmids were mobilizable but the 2337bp plasmids were non-mobilizable. The reason for the maintenance or transfer of these small plasmids is unclear as they do not appear to provide any known service or function to the bacterial host.

### Ecologically Associated Plasmids

By analysing the plasmids present per *E. coli* isolate we identified which plasmids co-existed together. Four mega conjugative plasmids (224Kb, 139Kb, 102Kb and 96Kb) co-existed in the same *E. coli* isolate (Table 1 and Supplementary information). In addition, subsets of these plasmids existed together either as three plasmids in isolate 3, which lacked the 96Kb plasmid, but contained four additional small plasmids each less than 5Kb) or as in isolate 42 which contained three plasmids and lacked the 224Kb plasmid and isolate 43 was missing the 139Kb plasmids but neither isolate 42 nor 43 contained additional plasmids (Table 1). The plasmids contained the replicon groups IncH1 and 2 and RepA (224kb), IncFIB and FIC (139Kb), IncFII (102Kb) and IncFII and IncI1 (96Kb). There was very little overlap in the AMR gene content across the plasmids, only the ANT(3’’)-IIa and *tetA* were present in two plasmids (224Kb and 102Kb). All plasmids were selected on tetracycline. The two plasmids lacking tetracycline resistance genes were only able to survive in the *E. coli* due to the presence of those with tetracycline resistance genes. The non-tetracycline resistance plasmids contained either only *bla*_TEM-1_ or no AMR genes (96kb and 139Kb, respectively). While virulence genes were co-transferred with AMR genes they were not transferred on the same plasmid. The sample containing the 224Kb, 139Kb, 102Kb plasmids contained four additional small plasmids (2337, 3463, 4358 and 4927 bp). However, when the mega-plasmid set also included the 96Kb plasmid there were no small plasmids present.

**Table 1.**
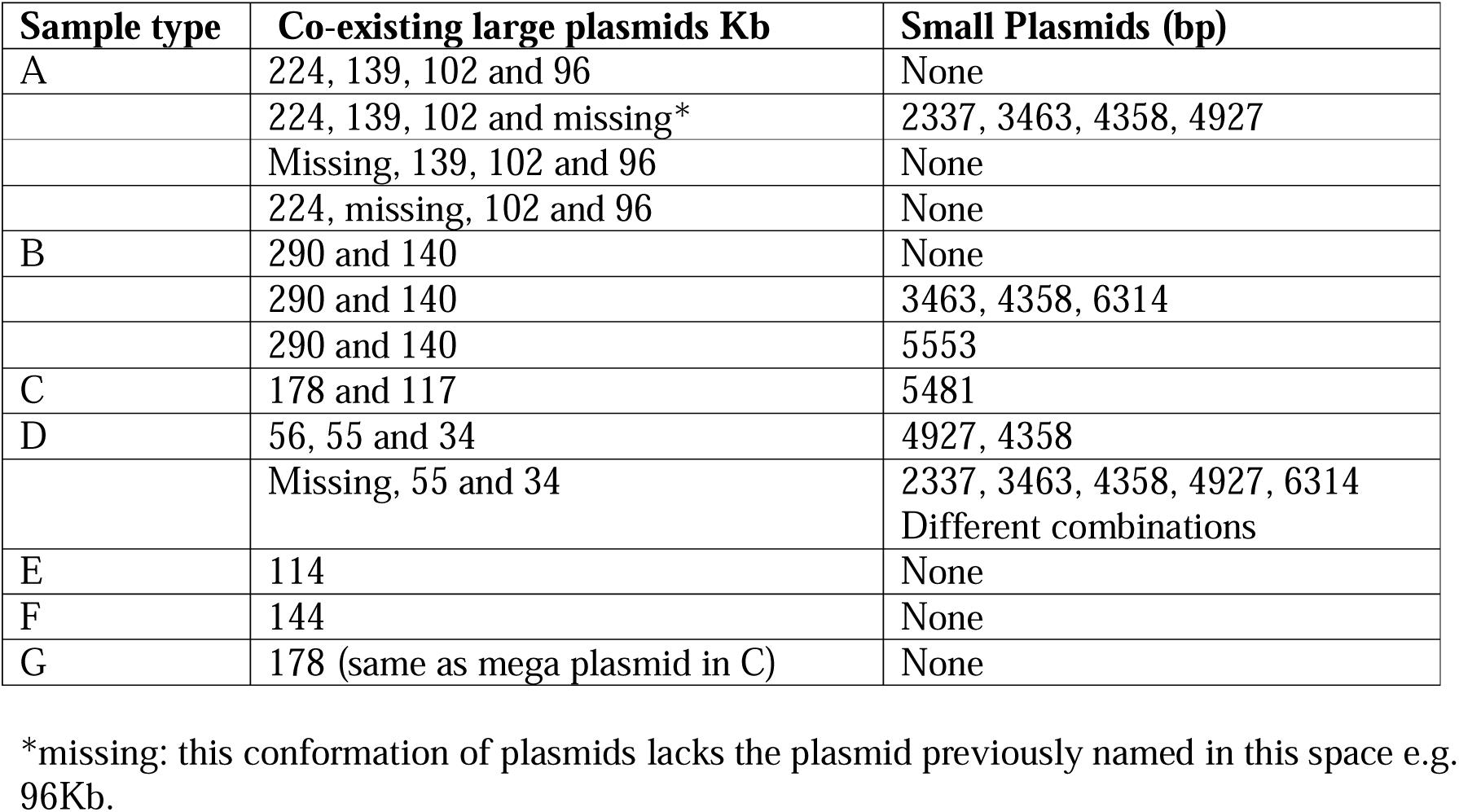
Co-existing plasmids within the same isolate.

There were two further sets of co-existing pairs of mega plasmids. One comprised 287/292Kb and 140Kb with or without small plasmids and the other comprised 178Kb and 117Kb with a 5.5Kb plasmid. The plasmids belonged to different replicon groups: 1) IncHI2A, IncH2B, RepA (287/292Kb) and IncFII (140Kb) and no replicon group (6Kb) and 2) IncIA/C2 (178Kb) and no identified replicon groups in 117Kb or 5.5Kb plasmids. The 287/292Kb plasmids isolated from *E. coli* selected on tetracycline carried 11 or 13 different AMR genes, including *tetD* conferring tetracycline resistance on which the colonies were selected, while the 140Kb plasmids contained no ARGs. Both the 287/292Kb and 140Kb plasmids contained mercury resistance genes and the 290KB plasmid additionally contained a further 17 metal resistance genes. The plasmids 287/292Kb and 140Kb co-existed with either a 5553bp plasmid, which was not detected in any other sample or with 3463, 4358, 6314bp plasmids together or with no small plasmids (Table 1). The co-existing 178Kb and 117Kb plasmids contained ARGs *ANT(3’’)-IIa*, *mphE*, *msrE* and *bla*_TEM-12_ (178Kb) and the 117Kb plasmid contained only *bla*_TEM-12_. *Escherichia coli* isolates containing these plasmids were selected on ampicillin, enabled by the *bla*_TEM-12_ genes. The 178Kb plasmid contained the *qacE* gene. The 178 – 117Kb *E. coli* isolates contained a 5481bp small plasmid, which was not detected in any other sample.

Medium sized plasmids (34, 55, 56Kb) were also detected with non-mega plasmids (Supplementary information). The *E. coli* containing these plasmids also contained up to five different small plasmids (Table 1) either all together in one isolate or a subset of the small plasmids across isolates. All of the detected small plasmids were also detected in the isolates containing mega plasmids, which suggests that the small plasmids are not aligning themselves with plasmids of a certain size nor selective trait. No isolates contained only small plasmids, even those that were mobilizable or contained the selective resistance gene.

### Ecologically Disassociated Plasmids

Some *E. coli* isolates only contained certain small plasmids in the absence of one of the large plasmids. *Escherichia coli* containing the 96Kb, 114Kb or 144Kb plasmids existed without small plasmids (Table 1). There was no pattern of a specific replicon group present on the plasmids that lacked small plasmids in comparison with those containing small plasmids. For example the replicon groups of mega plasmids transferred with the 3463bp plasmid were IncF, IncH and RepA (Supplementary information). The replicon groups of the 114Kb or 144Kb plasmids were IncF and the 96Kb plasmid contained IncF and IncI. There was no distinct pattern of gene content across the small plasmids to indicate why they were not co-transferred with the larger plasmids. There was also no distinct pattern across the 96Kb, 114Kb or 144Kb plasmids to indicate why the small plasmids were not transferred only in these cases. All were conjugative plasmids. The only unusual finding was the presence of three phages in the sample containing the 114Kb plasmids. If this is a phage plasmid this suggests that phage plasmids are not co-transferred with other plasmids. The 114Kb and 144Kb plasmids were the only plasmids detected solely as single plasmids in the samples.

### Genetic rearrangements of highly similar plasmids

Due to the methodology used we have selected groups of highly similar plasmids. When the plasmids were analysed they were not identical and many contained rearrangements.

### Small plasmid rearrangements

There were more rearrangements in the small plasmids than large plasmids. **Group 0** contained 18 plasmids all 4358bp in length. There were reshuffling and inversion events present across the plasmids. Within group 0 there were three basic sets of gene arrangements (Table 2A, B, C). The presence of *mbeD* in some versions of this plasmid are of interest as the TraM binds to one site at *oriT* of the ColE1 plasmid (28). Therefore, it is assumed that MbeD also binds to this site of ColE1. This means that MbeD plays a role in initiating plasmid transfer in the donor cells. **Group 1** comprised seven plasmids of 2337bp length. The rearrangement events were described as inversions and reshuffling events. Each plasmid consisted of an RNAi and two hypothetical genes (hypothetical gene 1 and hypothetical gene 2). However, while the RNAi and hypothetical gene one were the same length across all plasmids, there was variation in length of hypothetical gene 2 from 185bp to 542bp. Most plasmids contained a reshuffling event of approximately half the length of the plasmids in comparison with the others in the specific groupings above. The inversion events occurred across the entire plasmids and ranged in size from 180 bp to 2155 bp, which are the same inversions. **Group 2** comprised nine plasmids of 3463bp length and different gene combinations (Table 3). In four cases the *mobC* was replaced by a hypothetical gene and in one plasmid the lipoprotein gene was replaced by a hypothetical gene. *mobC* is a mobility gene, required for the transfer of the plasmid in the presence of a conjugative plasmid. **Group 3** comprised a group of 14 plasmids 4927bp in size (Table 4). Some plasmids were identical and contained no rearrangements (10D and 23D, 19D and 22B, 13D and 16D). The identical plasmids were isolated from different samples for example, 10D from WWTP timepoint 3, 23D from WWTP1 at timepoint 1. **Group 4** comprises 13 plasmids all 6314bp long with unaligned beginning/end, inversions and reshuffling events (Table 5). These small plasmids contain the *traD* gene. TraD is a hexameric ring ATPase that forms the cytoplasmic face of the conjugative pore and it interacts with TraM. Plasmid 12C and 25E lacked the *traD* gene.

**Table 2.** Plasmid rearrangements. x denotes presence of the gene, an absence of x indicates this gene is lacking and descriptions in the box correspond to the changes identified in this gene e.g. Hypo indicates that the gene is replaced by a hypothetical gene.

A Group 0 plasmids without additional genes inserted.
| Plasmid name | Antitoxin | *ccdB* | *mobA* | *mbeC* | *oriC* | hypo | rep protein | pseudoknot |
| --- | --- | --- | --- | --- | --- | --- | --- | --- |
| 10E | x | x | x | x | x | x | x | x |
| 12E | x | x | x | x | x | x |  | x |
| 13E | x | x | x | x | x | x | x | x |
| 14E | x | x | x | x | x | x |  | x |
| 15D | x | Hypo | x | x | x | x | x | x |
| 16E | x | x | x | x | x | x | x | x |
| 17E | x |  | x | x | x | x | x | x |
| 18D | x | x | x | x | x | x | x | x |
| 20E | x | x | x | x | x | x | x | x |
| 21C | x | x | x | x |  | x | x | x |
| 22C |  | x | x | x | x | x | x | x |
| 23E | x | x | x | x | x | x | x |  |
| 9E | x | x | x | x | x | x | x | x |

**Table 2 B. The rearrangement of Group 0 plasmids containing the *mbeD* gene.**
|  | Antitoxin | *ccdB* | *mbeD* | hypo | Hypo | Hypo | *mbeC* | *oriC* | Hypo | replication protein | pseudoknot |
| --- | --- | --- | --- | --- | --- | --- | --- | --- | --- | --- | --- |
| 11C | x | x | x | x | x | x | x | x | x | x | x |

**Table 2 C. The rearrangements of Group 0 plasmids containing the *mbeD* and *mobB* genes.**
|  | Antitoxin | *ccdB* | *mbeD* | *mobB* | Hypo | *mbeC* | *oriC* | Hypo | replication protein | pseudoknot |
| --- | --- | --- | --- | --- | --- | --- | --- | --- | --- | --- |
| 19E | x | x | x | x | x | x | x | x | x | x |
| 24C | x |  | x | x | x | x | x | x | x | x |
| 25A | x | x | x | x | x | x | x | x | x | x |
| 3B | x | x | x | x | x | x | x | x | x | x |

**Table 3:** Plasmid Group 2 rearrangements.

|  | Lipoprotein | Hypo | Putative mobilisation protein 3 | *mobC* | hypo | hypo |
| --- | --- | --- | --- | --- | --- | --- |
| 11D | x | x | x | x |  |  |
| 12F | x | x | x |  | x |  |
| 15E | x | x | x | x |  |  |
| 18E | x |  | x |  | x |  |
| 19F | x | x | x |  | x |  |
| 23F | x | x | x |  | x |  |
| 24D | x | x | x | x |  |  |
| 25B | x | x | x | x |  |  |
| 3C |  | x | x | x |  | x |

**Table 4:**
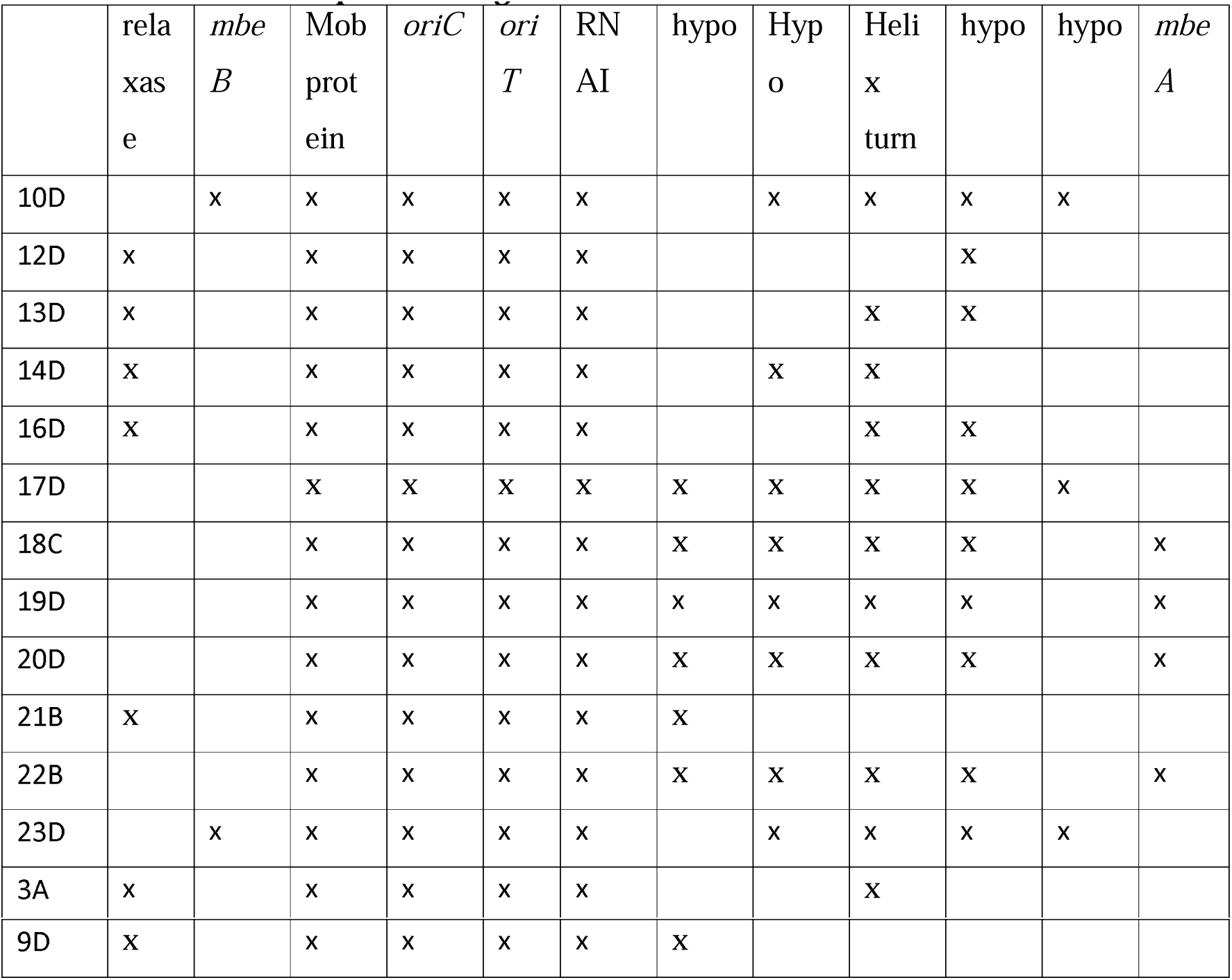
Plasmid Group 3 rearrangements.

**Table 5:**
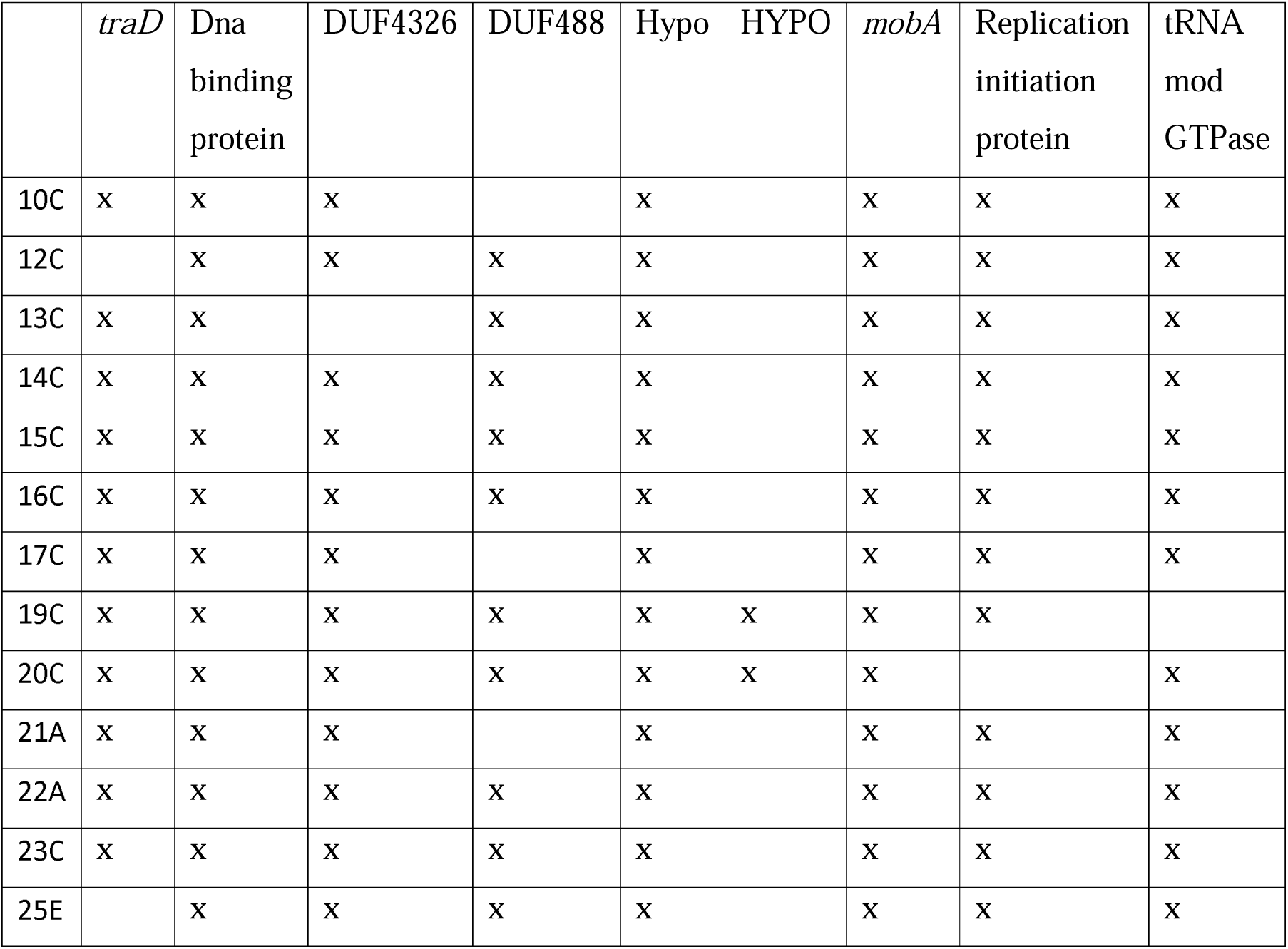
Plasmid Group 4 rearrangements.

|  | *traD* | Dna<br>binding<br>protein | DUF4326 | DUF488 | Hypo | HYPO | *mobA* | Replication<br>initiation<br>protein | tRNA<br>mod<br>GTPase |
| --- | --- | --- | --- | --- | --- | --- | --- | --- | --- |
| 10C | x | x | x |  | x |  | x | x | x |
| 12C |  | x | x | x | x |  | x | x | x |
| 13C | x | x |  | x | x |  | x | x | x |
| 14C | x | x | x | x | x |  | x | x | x |
| 15C | x | x | x | x | x |  | x | x | x |
| 16C | x | x | x | x | x |  | x | x | x |
| 17C | x | x | x |  | x |  | x | x | x |
| 19C | x | x | x | x | x | x | x | x |  |
| 20C | x | x | x | x | x | x | x |  | x |
| 21A | x | x | x |  | x |  | x | x | x |
| 22A | x | x | x | x | x |  | x | x | x |
| 23C | x | x | x | x | x |  | x | x | x |
| 25E |  | x | x | x | x |  | x | x | x |

### Rearrangements of Large plasmids

In theory, large plasmids have a comparatively smaller barrier to incorporating new genes when size is assumed to be broadly correlated with replicative burden (29). However, this was not the case in this study. The **Group 10** plasmids comprised 14 different plasmids ranging in size from 96358 bp to 97016 bp with the replicon groups IncI and IncFII. This is the only group of plasmids to contain the shufflon. Shufflons are present on IncI plasmids. The shufflon provides variety to the tip of the PilV adhesin. PilV recognises specific lipopolysaccharide structures on the surface of the recipient cell during mating (30). Within this group of plasmids rearrangements or changes occurred only within the shufflon region between 78901 and 80359bp across all plasmids. The event length ranged from 12 bp to 348bp deletions or insertions within this region (31). There were no other rearrangements on these plasmids. The conjugative plasmids in **group 7** and were 224Kb (n = 14), 287Kb (n = 2) or 292Kb (n = 5) in size. The plasmids of the same sizes were identical, resulting in three different plasmids in this group. In the 224Kb plasmids relocations with insertions were mainly based around IS elements or transposons that differ in the 224Kb group to the other two plasmids 287/292Kb. While they are all IncH plasmids with many conserved genes across the plasmids the variation is relatively large in the variable regions. For example a fragment of 11,214 bp was inserted into plasmids at the same location across the 287Kb or 292Kb plasmids in comparison with the 224Kb plasmids or in reverse a deletion occurred in the 224kb at location 23079bp. While in length plasmids 287 and 292kb were only 5kb in difference this is distributed across different sections of the plasmid via insertions and deletions. One example is a region of 4,952bp insert in 292Kb (229,318 to 234,270bp), which is absent from 287Kb. This region encodes an additional *sul1*, *hypA* (hydrogenase maturation factor), *ampR*, *qnrA* and an IS91 encoded by rPS27A. At the same location and prior to these genes both plasmids contain rPS27A. In a previous study the insertion of *qnrB* beside/near *oriC* resulted in increased bacterial mutation rate and was suggested that *qnrB* acts as a DNA replication initiation activator (30). However, there were no mutations in *oriC* genes of the plasmids in our study. **Group 5** comprises 16 plasmids. There were no statistical differences between 10A, 9B, 12A and 19A. There were also no changes between plasmids of the same sizes: 11A vs 24A, 20A, 13A, 16A, 21E, 22E, 23A and 14A, 15A, 17A. Thus we classified the 16 plasmids as five plasmids within this group. When plasmids 55291bp were compared with the 9B group there were collapsed tandem repeat regions. All events were within the origin of replication, which comprised collapsed repeats, tandem duplications, collapsed tandem repeats, deletion, insertion and duplication events. **Group 13** contains two identical plasmids: 46A and 47A which had no rearrangements. **Group 14** share a large proportion of genes that are not replicons or ARGs. They are dynamic in terms of One Health but have a close evolutionary history. Based on the identical nature of plasmids of the same size group 14 splits into 3 groups. 14i: 144,214bp (n = 2), 14ii: 138,989bp (n = 14), and 14iii: 101594bp (n = 15). Comparison across plasmid groupss e.g. 14i vs 14ii or 14iii demonstrated that they were as different from each other as a plasmid from another group with the same Inc group: 37A (144kb), 1B (IncFIB) (139Kb), 2C (102Kb). However, when all other proteins are considered, this isn’t the case. These plasmids are conserved in one sense but distinct in another - they are dynamic and conserved, which is rare. **Group 6** contains seven plasmids. They are identical to each other. These plasmids were present with other plasmids that contained rearrangements e.g. **group 7**. Group 7 plasmids had no AMR genes, mercury resistance genes and were 140,332bp. All plasmids contained insertion sequences and transposons but there were no variations within the plasmids. This suggests either a perfectly preserved operonic structure or they were very recently (but very rapidly) introduced prior to selection. There are 16 plasmids in **group 16**. There were no AMR, no metal resistance, no virulence genes present. They are all IncP plasmids. Plasmids of the same size were identical. This resulted in seven unique plasmids for rearrangement comparisons. All rearrangements within this group comprised tandem duplications or collapsed tandem repeats within the region of the origin of replication. The rearrangements varied in size from 50bp to 332bp. All tandem duplications ended at position 1745bp and varied in length from 145 bp to 264bp and all occurred in the origin of replication. No rearrangements were detected in the other regions of these plasmids.

When analysing the rearrangements of large plasmids three patterns emerged:

1. All plasmids within a groups of IncF plasmids or IncA/C were identical i.e. no rearrangements occurred.
2. Plasmids contained rearrangements close to or within the origin of replication in IncH or IncN or IncP plasmids
3. Plasmids contained rearrangements within the shufflon on the IncI plasmids

## Discussion

Analysis of AMR across One Health has often been discussed using culture of pathogens, or DNA based techniques, which identify the presence of AMR genes but rarely the circularised plasmids containing the ARGs. Co-plasmid existence is a little studied area of plasmid ecology and dynamics. In most studies the analysis has been to investigate the plasmid pairs in terms of costs or benefits, the ability to exclude a plasmid or the potential for one plasmid to provide a benefit to the other *in vitro*.

Understanding the ecology and evolution of plasmids and especially AMR plasmids is vital to enable us to understand their transmission across One Health. The plasmid paradox suggests that theoretically plasmids should not exist as the costs associated with plasmid maintenance would result in non-beneficial plasmids being removed by negative selection, whereas the genes present under positive selection should be captured to the bacterial chromosome, which would result in loss of the redundant plasmid (6). This assumes that the entropic fate of plasmids is to become chromosomal. However, we know that this does not happen due to the wide array of plasmids present in nature. Solutions to this plasmid paradox have suggested ecological and evolutionary reasons for the existence of plasmids (6). These solutions are based on *in vitro* studies of plasmids transferred into one or two bacterial species. One study demonstrated positive epistasis between small and large plasmids conferring resistance to antibiotics and a second study, while using different methods, showed a synergistic effect of two large conjugative plasmids co-infecting the same species (3, 33). Our study provides evidence to show that plasmid co-existence occurs widely in a complex biome and that there is some force that shapes the plasmid community content.

Our data has demonstrated:

1. Large conjugative plasmids with selective advantages allow large conjugative plasmids with the same or no selective advantage to co-exist and benefit from their selective advantage.
2. Large conjugative plasmids co-exist with small mobilizable plasmids with no apparent benefit to the large plasmid but the small plasmids gain promiscuity.
3. Small plasmids were not detected in isolation and therefore must have a co-dependency on the large plasmids.
4. The pattern of small and large co-existing plasmids suggests a selective exclusion property of large plasmids for small plasmids.
5. The small plasmids were highly rearranged indicating that they are more fluid DNA entities than the large plasmids suggesting either a greater variety of highly similar small plasmids exist in nature or that they are capable of many rearrangements.

Applying the current solutions of the plasmid paradox to these findings we discuss how these solutions relate to the communities of plasmids in this study. Our results suggest that positive epistasis promotes plasmid persistence in WWTP populations in a similar manner to that identified in pure bacterial populations (3).

### Ecological solution 1: infectious transmission

While infectious transmission through conjugation relates only to conjugative plasmids we will use the same metrics for all our data. This concept states that plasmids survive at population level by horizontal transmission as an infectious element. This implies a parasitic symbiotic relationship. Based on what we observed here, plasmids were much less parasitic than would be observed in a phage-cell ecosystem. We suggest this is a subcellular mixed “species” flocking relationship observed between plasmids and bacteria. While this would increase the potential fitness cost it results in a wider array of selective advantages conferred to the individual plasmids when in a community rather than single entities. This is displayed in the communities of varying large plasmids of 224Kb, 139Kb, 102Kb and 96Kb. This mode of transmission enables plasmids to hide until their selective advantage is required by utilising the selective advantage of the other plasmid(s). These large conjugative plasmids communities may also include small plasmids, that vary in their combinations. Such variety in plasmid communities enables the bacterial host to be ready for many selective environments and thus enhance their survival. It also enables the persistence of small mobilizable but non-conjugative plasmids.

### 2. Plasmid properties vary across host backgrounds

As we have investigated the plasmids in *E. coli* we do not know the variation of properties across host backgrounds. However, others have identified that the presence of two plasmids with the same trait confer an additive phenotypic effect rather than of one or the other. In addition, the communities varied in their arrangements of plasmids e.g. containing small plasmids with some and lacking small plasmids with other. In addition, the levels of rearrangements of the plasmids in the plasmid communities demonstrates a plasmid variation across the total plasmid communities. Thus it is not only the plasmid host that impact plasmid properties but also their plasmid communities content.

### Solution 3: Interactions with other plasmids

While plasmid persistence has been demonstrated to be effected by other plasmids these experiments have analysed *in vitro* plasmid combinations and were limited to two plasmids, most frequently one large and one small plasmid. Our study suggests that synergistic epistasis is more common than previously thought as most plasmids in our study co-existed with at least one other and up to four other plasmids. Further experimental analysis across these plasmids is required to test the importance of plasmid co-existence in the proliferation and persistence of plasmids as communities.

### Solution 4: Source-sink spill over transmission

This solution suggest that plasmids can exist in a microbiome by persisting in a species more proficient at acting as a host for that plasmid and can then spill over into the less proficient host (6). A plasmid went extinct in one species in the absence of selective pressure but was capable of residing in another species and then reinfecting the original species. In our study we suggest that plasmid communities act as a source of a spill over plasmid transmission. Our data contained several examples where one plasmid went extinct. However, there were other *E. coli* that contained the same plasmid in another configuration i.e. persisted. We suggest that communities of plasmids act as independent, or semi-independent, communities and configure themselves in different combinations in order to preserve the total plasmid community but without the need to preserve all plasmids in the same *E. coli* concurrently. The population is ready for many selective pressures but individual *E. coli* do not carry the burden of all plasmids.

Compensatory mutations in the plasmid or chromosome have provided a reduced fitness cost of carrying the plasmid to the host bacteria. One compensatory mutation study showed that increased plasmid stability in *E. coli* was due to large deletions of approximately 25Kb when in *E. coli* but never in *Klebsiella pneumoniae* (34). Within our study we identified genetic rearrangements across the small plasmids at a higher frequency than the large plasmids. This suggests that small plasmids are more plastic and readily capable of change. It also results in a wider range of variable small plasmids within the plasmid communities of the population. Currently, we do not know how these rearrangements benefit the plasmid community. Large plasmids contained more restricted rearrangements with many around the region of the origin of replication. This suggests that the large plasmids are not evolving as quickly as the small plasmids and are perhaps slower at adapting to change. We have not analysed the bacterial host chromosome. It will be interesting to analyse the impacts of these various rearrangements on the plasmid community stability and maintenance over time and across selective pressures.

## Conclusions

In conclusion we propose that plasmid co-existence is a very important factor to understand the plasmid dynamics and ecology in nature and how those plasmids generate small changes to their genetic content enables them to evolve and maintain a variety of highly similar but different plasmids within a bacterial population. This results in changing how we study plasmids from the impact of single plasmids on the host bacteria or community to the interactive and community of plasmids capable of co-existing in different combinations within either a bacterial species, as demonstrated in this study, or within a population of bacteria.

## Acknowledgements

This work was funded by co-funded PhD studentship from The Kathleen Lonsdale Institute for Human Health Research and Environmental Protection Agency (EPA) to Dr Cian Smyth and Prof Fiona Walsh. The JPI Water Stare project was funded by the EPA under the EU JPI Water programme.

## Author contributions

CS performed the experiments, data analysis, RL performed data analysis, TTD performed experiments, TTD and FW generated the research project, FW supervised the work, obtained the funding and wrote the manuscript.

## Data availability statement

Data is available in NCBI via BankIT under the accession number XYZ

